# RNA components of the spliceosome regulate tissue- and cancer-specific alternative splicing

**DOI:** 10.1101/326983

**Authors:** Heidi Dvinge, Jamie Guenthoer, Peggy L. Porter, Robert K. Bradley

## Abstract

Alternative splicing of pre-mRNAs plays a pivotal role during the establishment and maintenance of human cell types. Characterizing the *trans*-acting regulatory proteins that control alternative splicing in both healthy and malignant cells has therefore been the focus of much research. Recent work has established that even core protein components of the spliceosome, which are required for splicing to proceed, can nonetheless contribute to splicing regulation by modulating splice site choice. We here demonstrate that the RNA components of the spliceosome likewise influence alternative splicing decisions and contribute to the establishment of global splicing programs. Although these small nuclear RNAs (snRNAs), termed U1, U2, U4, U5, and U6 snRNA, are present in equal stoichiometry within the spliceosome, we found that their relative levels vary by an order of magnitude during development, across tissues, and between normal and malignant cells. Physiologically relevant perturbation of individual snRNAs drove widespread gene-specific differences in alternative splicing, but not transcriptome-wide splicing failure. Genes that were particularly sensitive to variations in snRNA abundance in a breast cancer cell line model were likewise preferentially mis-spliced within a clinically diverse cohort of invasive breast ductal carcinomas. As aberrant mRNA splicing is prevalent in many solid and liquid tumors, we propose that a full understanding of dysregulated pre-mRNA processing in cancers requires study of the RNA as well as protein components of the splicing machinery.

## INTRODUCTION

Alternative pre-mRNA splicing, which permits the expression of multiple transcript isoforms from a single gene, affects almost all multi-exon human genes (Wang et al. 2008). Alternative splicing plays correspondingly crucial roles during normal biological processes such as development and cell type specification (Graveley 2001; Pan et al. 2008; Yang et al. 2016; Ellis et al. 2012; Kalsotra and Cooper 2011; Chen and Manley 2009). Conversely, dysregulation of alternative splicing characterizes many genetic diseases and cancers (Dvinge and Bradley 2015; Climente-González et al. 2017; Dvinge et al. 2016; Scotti and Swanson 2016), and is sufficient to drive disease initiation, progression, and therapeutic response (Quesada et al. 2011; Zhang and Manley 2013; David and Manley 2010; Yoshida et al. 2011; Graubert et al. 2011; Papaemmanuil et al. 2011; Imielinski et al. 2012; Martin et al. 2013). Accordingly, substantial effort has been devoted to identifying and characterizing the factors that control alternative splicing programs in both healthy and diseased cells.

Factors involved in the splicing process can be roughly categorized as “basal” or “regulatory,” depending upon whether or not they are required for all splicing. In this simplified view, basal factors are required to catalyze the splicing process itself, while regulatory factors promote or repress splicing. Canonical basal factors include those that constitute the spliceosome, whose core components are the NineTeen Complex (NTC) and five small nuclear ribonucleoprotein complexes (snRNPs, pronounced ‘snurps’), known as U1, U2, U4, U5, and U6 (**Fig. 1A**). Canonical regulatory factors, on the other hand, may not be part of the core spliceosome itself. Instead, these regulatory factors typically bind specific enhancer or silencer sequences in pre-mRNA to promote or repress splicing (Fu and Ares 2014; Gerstberger et al. 2014). Canonical basal factors are ubiquitously expressed in all cells since they are required for splicing to occur, while cell type-specific expression of regulatory factors contributes to the establishment and maintenance of distinct splicing programs in different cells. Dysregulated expression or mutations affecting specific splicing factors can alter the normal splicing program to drive genetic, dysplastic, and neoplastic disease (Dvinge et al. 2016; Raj and Blencowe 2015).

**Figure 1:**
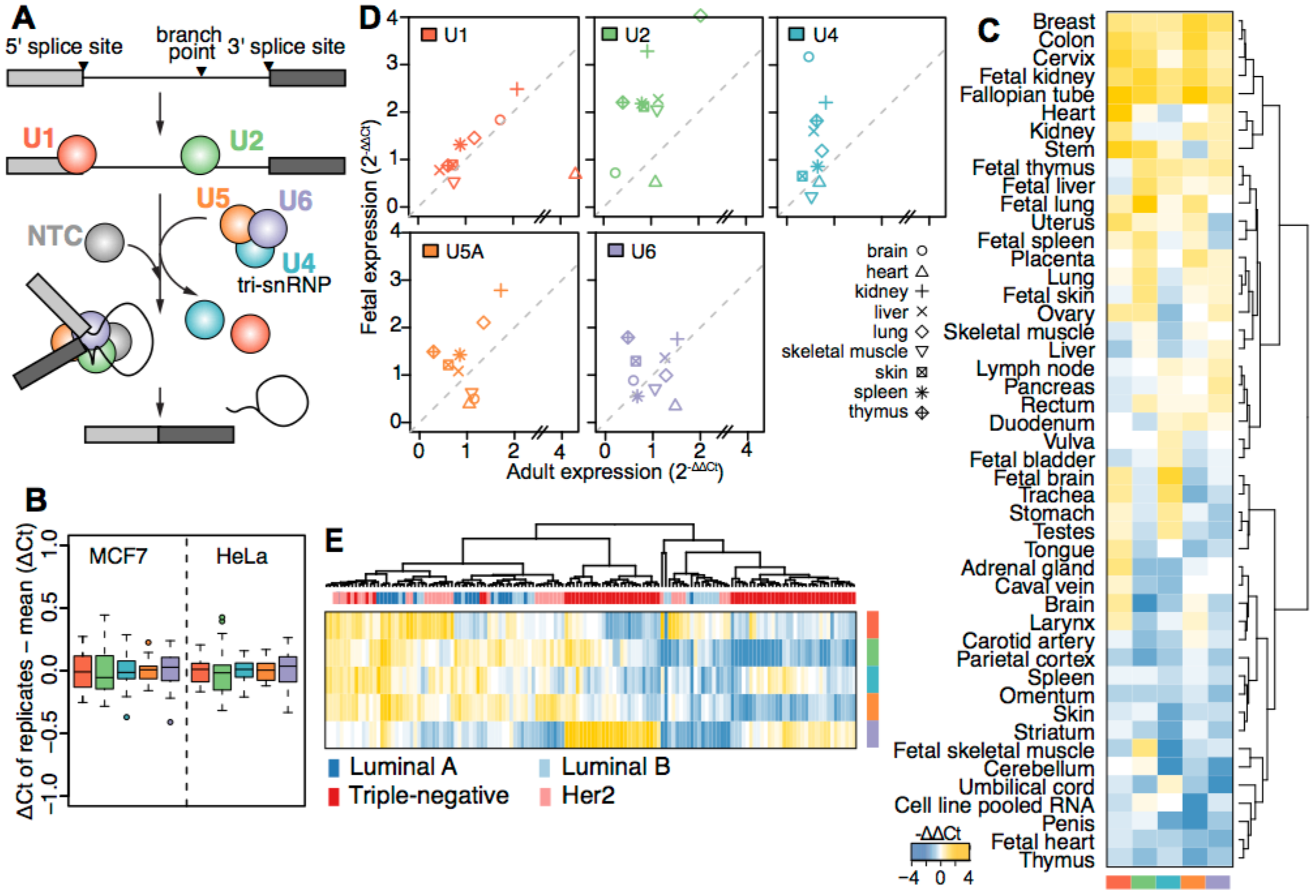
Spliceosomal snRNA abundance is highly variable. **(A)** Simplified schematic of a single round of splicing, showing individual steps: recognition of the 5’ and 3’ splice sites by the small nuclear ribonucleoprotein complexes (snRNPs) containing the U1 and U2 snRNAs, respectively; recruitment of the U4/U6.U5 tri-snRNP; exit of the U1 and U4 snRNAs and rearrangements of the snRNPs into the conformation required for the active spliceosome; excision of the intron lariat and ligation of the two adjacent exons. **(B)** Reproducibility of ΔCt values from our microfluidic real-time quantitative PCR-based assay to measure snRNA levels across five biological and three technical replicates, using the MCF7 and HeLa cell lines. For the calculation of ΔCt, the median of RN7SK, RN7SL1 and 5S rRNA within each tissue was used as a reference. **(C)** Heatmap of relative snRNA abundance across 36 healthy tissues and pooled cell line RNA. ΔCt values calculated as in (B). ΔΔCt values are relative to the median values across all tissues. Yellow; lower ΔΔCt (*i.e.*, higher expression), blue; higher ΔΔCt. **(D)** Expression level (2^−ΔΔCt^) of snRNAs in adult versus fetal samples from identical tissues. Colors as in (A). **(E)** Variations in snRNA abundance across 144 primary breast cancer specimens, calculated as in (C). The column color bar indicates the intrinsic breast cancer subtypes, as defined by immunohistochemistry (IHC) on these samples.

While attractive in its simplicity, recent studies have demonstrated that categorizing factors as purely basal or purely regulatory is inaccurate. Core components of the spliceosome can play regulatory, in addition to basal, roles. Even if a particular core spliceosomal protein is required for splicing to occur, developmental stage- or tissue-specific variation in its expression level can confer a regulatory role on the protein. For example, the core spliceosomal protein SmB/B’ regulates splicing of a cassette exon within its own pre-mRNA, as well as hundreds of other cassette exons (Saltzman et al. 2011). Other core spliceosomal proteins participate in regulation of alternative splicing (Perez-Santángelo et al. 2014; Wickramasinghe et al. 2015; Pacheco et al. 2006; Park et al. 2004) and display tissue- or development-specific expression patterns (Grosso et al. 2008). These studies suggest that the repertoire of splicing factors that play regulatory roles may be substantially larger than is currently realized.

Since core protein components of the spliceosome can act as regulatory factors, we wondered whether the RNA components of the spliceosome might similarly contribute to splicing regulation. Each of the five snRNPs contains a cognate U-rich small nuclear RNA (snRNA). The U1 and U2 snRNAs are responsible for recognizing the 5’ splice site and branch point upstream of the 3’ splice site, followed by the U4/U6.U5 tri-snRNP joining the spliceosome prior to the rearrangements that ultimately leads to the U6 snRNA catalyzing the actual splicing reaction (Fica et al. 2013). These snRNAs are present in constant stoichiometry within the spliceosome and are strictly required for splicing to occur. snRNAs therefore seem like canonical basal factors whose depletion would simply lead to global reductions in splicing efficiency. Instead, however, several disease-associated perturbations in snRNA levels give rise to cell type-specific changes in splicing that preferentially affect specific genes (Zhang et al. 2008; Ishihara et al. 2013; Jia et al. 2012).

By analogy with snRNA perturbation in disease, endogenous variation in snRNA levels could potentially enable these RNAs to regulate alternative splicing. However, it is unknown whether such variation occurs in healthy cells. Only 100 to 200 nt in length, snRNAs are not detected by most large-scale assays routinely employed in functional genomics (*e.g.*, microarrays and most RNA-seq protocols), unless those assays are specifically designed to target short non-polyadenylated RNA species. snRNAs levels could potentially vary during development, between cell types, or in healthy versus cancerous cells, but their levels have never been systematically quantified across those biological axes.

Here, we systematically tested the hypothesis that endogenous variation in snRNA levels confers regulatory capacity on these RNAs. We discovered that the relative abundance of different snRNAs varies by an order of magnitude during development, across tissues, and between healthy and malignant cells. Mimicking this physiological variability by ectopically perturbing snRNA expression levels drove gene-specific changes in alternative splicing, but not widespread splicing failure. Genes that were particularly sensitive to snRNA level perturbation in cultured breast epithelial cells were preferentially mis-spliced within a breast cancer cohort. Our results demonstrate that unexpected variability in snRNA levels contributes to the establishment of global splicing programs in both healthy and diseased cells.

## RESULTS

### Absolute and relative snRNA abundance exhibit extreme variation

We first tested whether snRNA levels were relatively constant, as might be expected from their equal stoichiometry within the spliceosome itself, or if instead snRNA levels were variable, as is common for regulatory splicing factors. We used a microfluidic platform to develop a high-throughput quantitative real-time PCR assay to measure levels of all five snRNAs. As U5 has five distinct sequence variants (U5A, U5B, U5D, U5E, and U5F), we focused on the most abundant form, U5A (Sontheimer and Steitz 1992; Krol et al. 1981). We confirmed the robustness and reproducibility of our microfluidic assay by measuring snRNA levels across five biological and three technical replicates from two distinct cell lines (MCF7 and HeLa), with average Ct values of 10.38 ± 0.18 across replicates (**Fig. 1B**).

We used our high-throughput assay to systematically measure snRNA abundance across three distinct biological axes for which splicing is known to play critical roles: between tissues, during development, and in healthy versus cancerous cells. We quantified snRNA levels across diverse tissues derived from healthy donors, including 36 adult tissues and 10 fetal tissues, as well as across a cohort of 144 primary breast cancer specimens. We observed an unexpected and striking degree of variability in both absolute expression levels and relative expression levels of each snRNA across all three biological axes. Within a given tissue, different snRNAs were expressed at levels varying by up to eight-fold with respect to each other; conversely, each snRNA exhibited a similar degree of expression variability across different tissues (**Fig. 1C**). U1 was present in excess of other snRNAs, as expected from previous studies (Baserga and Steitz 1993). None of the snRNAs exhibited coordinated expression levels, including U4, U5A, and U6, even though they play intertwined roles as components of the U4/U6.U5 tri-snRNP.

We next compared snRNAs levels between the nine tissues for which both fetal and adult samples were available. Relative levels of U1 were very similar between fetal and adult tissues, except for heart, where adult expression was much higher. In contrast, U2 and U4 were expressed much more highly in fetal relative to adult samples. Levels of U5A and U6 were not consistently lower or higher in either development stage (**Fig. 1D**).

Finally, we compared relative snRNA abundance across a cohort of 144 invasive breast cancer samples. Biopsies were selected to represent all breast cancer subtypes (Sorlie et al. 2001), with a focus on the aggressive triple-negative tumors (N = 66 triple-negative; 22 Luminal A; 22 Luminal B; 34 Her2 positive). Even though all samples were taken from breast ductal carcinoma, they exhibited a similar degree of variability in relative snRNA levels as we observed across our entire panel of human tissues. This variability in snRNAs levels was not random. An unsupervised cluster analysis of the cohort, based solely upon snRNA levels, revealed that most samples exhibited subtype-specific patterns of snRNA expression (**Fig. 1E**). Interestingly, triple-negative samples clustered into two distinct groups with different patterns of relative snRNA expression, perhaps reflecting the well-known heterogeneity of this subtype (Lehmann et al. 2011). We conclude that snRNA levels are extremely variable across a wide range of biological conditions.

### Physiological perturbation of snRNA levels modulates alternative splicing

We next sought to test whether the high physiological variability in snRNA levels that we observed might contribute to the establishment of global splicing programs. Since the abundance of individual snRNAs was not coordinately regulated across tissues or cancer biopsies, we hypothesized that perturbing the expression of a specific snRNA within physiological ranges would modulate splicing. Short nuclear non-coding RNAs are not amenable to RNAi (Ploner et al. 2009) and most snRNAs are present in the genome as multi-copy genes, rendering genetic knockouts infeasible. We therefore transiently transfected cells with chemically modified antisense oligos to trigger RNaseH-mediated degradation of each specific snRNA, a strategy which has proven effective for targeting U1 snRNA (Liang et al. 2011; Vickers et al. 2011). We focused our knockdown (KD) studies on U1, U2, U4 and U6, but not U5, since its distinct sequence variants makes it resistant to efficient targeting by our antisense oligo strategy.

We efficiently depleted each of U1, U2, U4, and U6 snRNAs to levels that were comparable to the variation observed across healthy tissues and our breast cancer cohort in MCF7 and HeLa cells (**Fig. 2A**). We performed matched RNA-seq following depletion of each snRNA or transfection with a non-targeting control oligo. We quantified genome-wide alternative splicing of competing 5’ and 3’ splice sites, retained introns, and cassette exons, removal of constitutive introns, and alternative splicing of constitutive splice junctions as previously described (Dvinge and Bradley 2015) (Supplementary Table 1).

**Figure 2:**
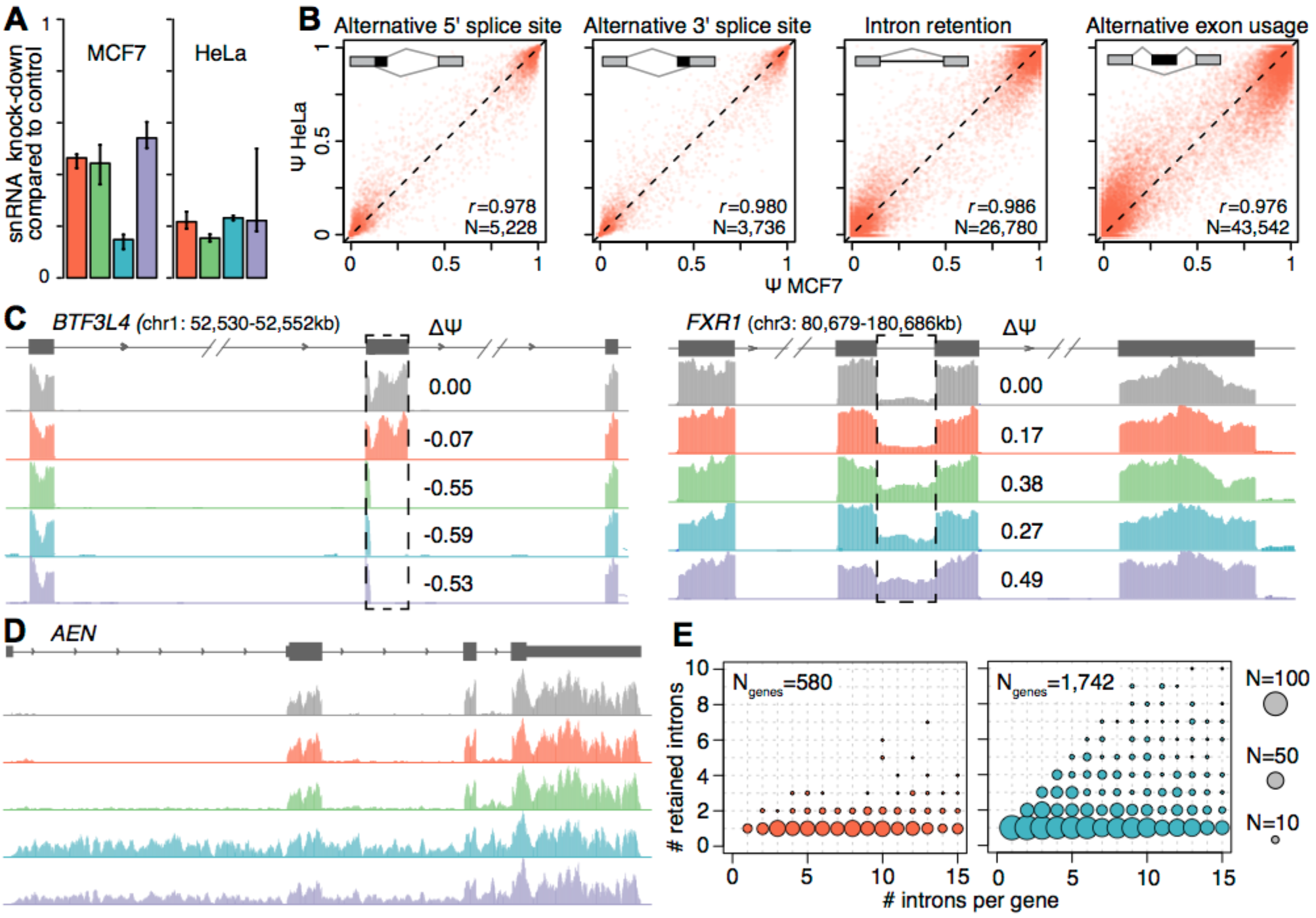
Physiological depletion of snRNAs modulates alternative splicing. **(A)** Levels of each snRNA following knock down (KD) in MCF7 and HeLa cells. Levels are relative to cells transfected with a scrambled control oligo. Error bars illustrate 95% confidence intervals, calculated using the balanced repeated replication technique across three technical replicates of each measurement. **(B)** Alternative splicing induced by U1 snRNA KD in MCF7 versus HeLa cells. Each dot illustrates an individual splicing event (alternative 5’ splice sites, alternative 3’ splice sites, exons, and introns), represented as percent-spliced-in (PSI, Ψ) values. Plot restricted to events with at least 20 informative reads. **(C)** Alternative splicing of an exon within Basic Transcription Factor 3 Like 4 (*BTF3L4*, left) and an intron within FMR1 Autosomal Homolog 1 (*FXR1*, right). Dashed boxes indicate the differentially spliced regions. ΔΨis defined as the fraction of transcripts containing the alternatively spliced region in snRNA KD versus control KD samples. **(D)** Intron retention across an entire transcript of Apoptosis Enhancing Nuclease (*AEN)* following U4 or U6 snRNA KD. Number of introns per gene (x axis; plot restricted to genes with <= 15 introns) versus introns with a statistically significant increase in intron retention (y axis) following KD of U1 (left) or U4 (right) snRNA. Circle size is proportional to the number of genes for each (x,y) combination.

We observed concordant patterns of global splicing changes in MCF7 and HeLa cells following KD of each snRNA relative to control KD samples. Changes in isoform usage for alternative 5’ splice site, alternative 3’ splice site, intron retention, and cassette exon events each exhibited Pearson correlations >0.97 between MCF7 and HeLa cells for each snRNA (**Fig. 2B**, illustrated for U1 KD samples). Given the highly concordant nature of these changes in splicing, we therefore focused subsequent analyses on data from MCF7 cells. This cell line was established from a pleural effusion of a breast adenocarcinoma patient, and it is therefore particularly biologically relevant to our breast cancer cohort.

Having established our model system, we next tested whether snRNA KD resulted in globally inefficient or failed splicing (suggestive of a purely basal role for snRNAs) or instead preferentially affected specific splice sites, exons, or introns (suggestive of a potential regulatory role for snRNAs). We did not observe classic hallmarks of globally inefficient splicing, such as widespread intron retention. Instead, most splicing changes affected single exons in an snRNA-dependent manner, while adjacent upstream and downstream exons were recognized with apparently normal efficiency (**Fig. 2C**, left). We did observe intron retention, suggestive of failure to recognize splice sites or catalyze splicing; however, typically only single introns were affected while neighboring introns within the same transcript were spliced efficiently (**Fig. 2C**, right).

While most genes contained only a single exon or intron that was affected by snRNA KD, a small subset of genes were strikingly sensitive to perturbation of snRNA levels. For genes such as *AEN*, we observed increased intron retention for most or all introns within the gene following U4 or U6 KD (**Fig. 2D**). We quantified this effect genome-wide by enumerating the number of retained introns relative to the total number of introns for each gene. For U1 KD, almost all affected genes contained just one retained intron, regardless of the gene structure (**Fig. 2E**, left). For U4 KD, in contrast, many genes contained multiple retained introns, and a small number of genes exhibited complete intron retention (**Fig. 2E**, right). The consequences of U2 and U6 KD were similar to those observed for U1 and U4 KD, respectively.

### Different snRNAs have consistent roles in constitutive and alternative splicing

As each snRNA has a distinct and well-characterized role in splicing, we wondered whether the observed differential splicing following KD of each snRNA was consistent with the stage of the splicing process when each snRNP joins the spliceosome (**Fig. 1A**). We clustered control, U1, U2, U4, and U6 KD samples according to how each affected alternative splicing of competing 5’ and 3’ splice sites, intron retention, and alternative exon usage (**Fig. 3A**). U1 and U2 KD primarily impacted splice site recognition and cassette exon inclusion, consistent with their respective roles in binding the 5’ splice site and branchpoint upstream of the 3’ splice site (**Fig. 3A-B**). U4 and U6 snRNAs, in contrast, primarily affected intron retention, consistent with their roles in splicing catalysis following splice site recognition (**Fig. 3A,C**).

**Figure 3:**
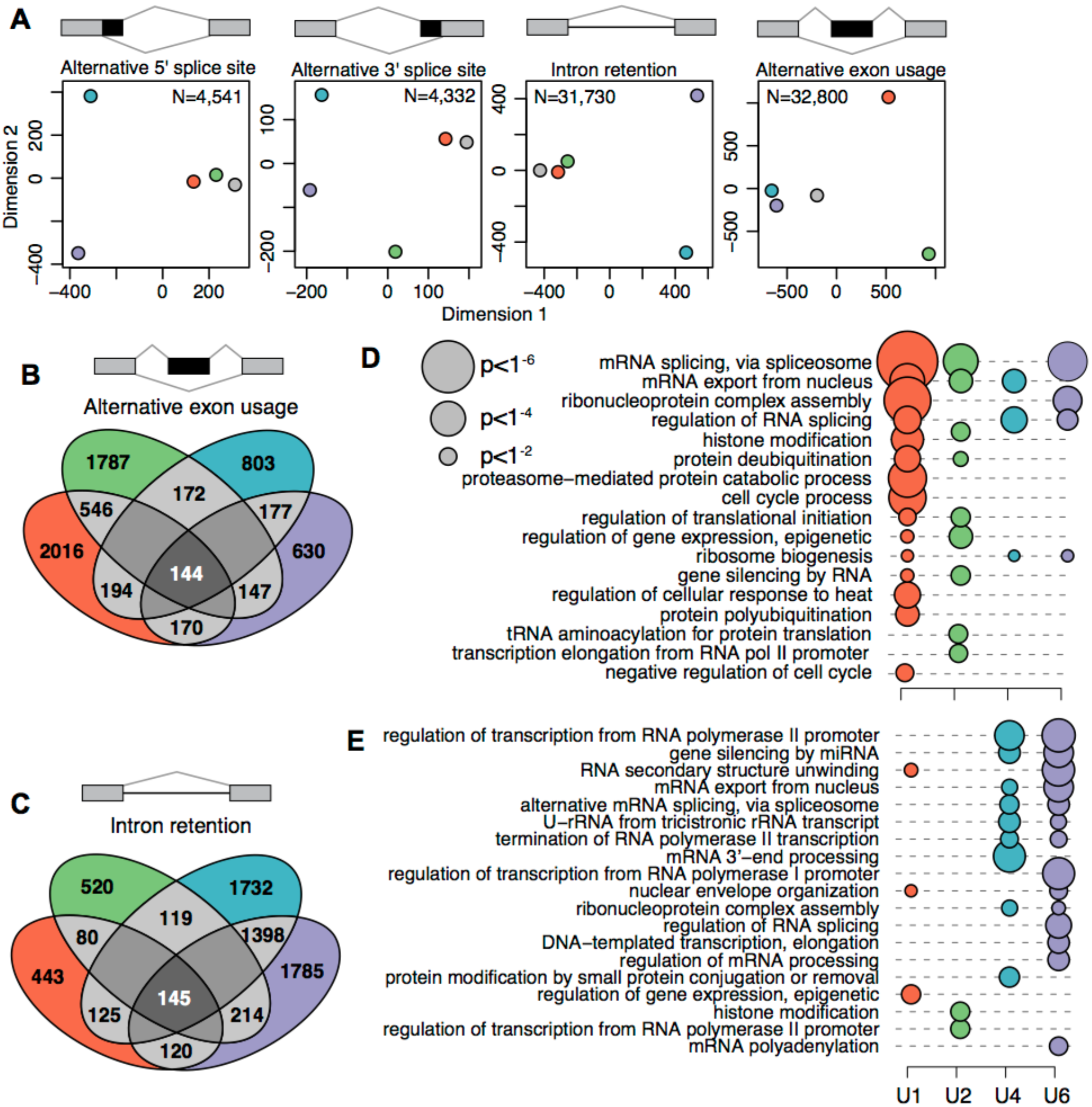
The functional of individual snRNAs is similar in basal and alternative splicing. **(A)** Two-dimensional clustering (multidimensional scaling) of samples based on transcriptome-wide splicing of different categories of splicing events, as in Fig. 2B. The clustering was based on the events with the largest degree of variation across samples, as indicated by N. Colors as in Fig. 1A. **(B)** Overlap between exons and **(C)** introns that are differentially spliced following KD of each snRNA in MCF7 cells. Most events are unique to depletion of individual snRNA. **(D)** Enrichment of Gene Ontology (GO) biological process categories for genes containing differentially spliced exons or **(E)** introns. The displayed terms were selected based on their combined significance across the four samples. Only terms with at least two ancestors are illustrated. Circle size is proportional to the statistical significance.

### snRNA KD has distinct but convergent biological consequences

We next tested whether the splicing changes that we observed were specific to particular snRNAs, or if instead perturbation of any snRNA preferentially affected the same transcripts. Approximately 27% and 29% of differentially spliced cassette exons were shared between U1 and U2 KD, while 52% and 50% of differentially retained introns were shared between U4 and U6 KD. This overlap is likely due to the related roles of U1 and U2 snRNA in defining splice sites and U4 and U6 KD in later stages of spliceosome formation and subsequent splicing catalysis. Nonetheless, perturbation of each snRNA induced a largely distinct program of alternative splicing (**Fig. 3B-C**). We conclude that a small subset of the transcriptome is sensitive to perturbation of any snRNA, but that most transcripts respond to inhibition of only a single snRNA, at least within physiological ranges of snRNA KD.

Since each snRNA was associated with both shared and distinct differential splicing following KD, we wondered whether the same would hold true for the downstream biological pathways affected by those splicing changes. We used Gene Ontology analysis to identify pathways that were enriched for genes containing differentially spliced exons or retained introns, as those were the predominant classes of differential splicing. Many snRNA-modulated exons were located within genes encoding proteins involved in RNA processing, including splicing as well as mRNA transport (**Fig. 3D**). Protein metabolism was also affected by differential splicing, through regulation of translation as well as protein stability in the form of ubiquitination. Retained introns were enriched within transcripts involved in post-transcriptional control of RNA processing or localization, but transcription itself was also over-represented (**Fig. 3E**).

snRNA-mediated differential splicing affected each of the highlighted biological processes by altering transcripts expressed by multiple genes within each pathway. For example, for mRNA export from the nucleus, altered exon inclusion affected multiple components of the nuclear pore complex (*e.g., NUP160, NUP188, NUP210, NUP85, NUP98*) as well as the TREX (TRanscription-EXport) complex (*e.g., CHTOP, NFX1, THOC2, THOC6*). Protein stability was affected via differential splicing of cassette exons within genes encoding F-box proteins (*e.g., FBXO22, FBXO31, FBXO4, FBXO44, FBXO7, FBXO9*) as well as genes encoding E2 ubiquitin-conjugating enzymes (*e.g., UBE2B, UBE2G1, UBE2G2, UBE2N, UBE2Q2, UBE2R2*).

### snRNA-mediated alternative splicing is affected by mechanisms acting in *cis* as well as *trans*

We next sought to determine why some, but not most, splice sites were preferentially sensitive to variable snRNA levels. As snRNA KD depletes levels of factors that are required for splice site recognition or splicing catalysis, we hypothesized that affected and unaffected splice sites might be particularly “weak” or “strong,” respectively. We therefore measured the approximate strength of each 5’ and 3’ splice site by comparing it to the genome-wide consensus for those splice sites using the MaxEnt method (Yeo and Burge 2004). Surprisingly, we observed the opposite pattern than what we hypothesized. Cassette exons that were preferentially included following KD of any snRNA had much weaker 5’ and 3’ splice sites than the genomic average, while cassette exons that were preferentially excluded following U1 or U2 KD had modestly stronger 5’ and 3’ splice sites (**Fig. 4A**).

**Figure 4:**
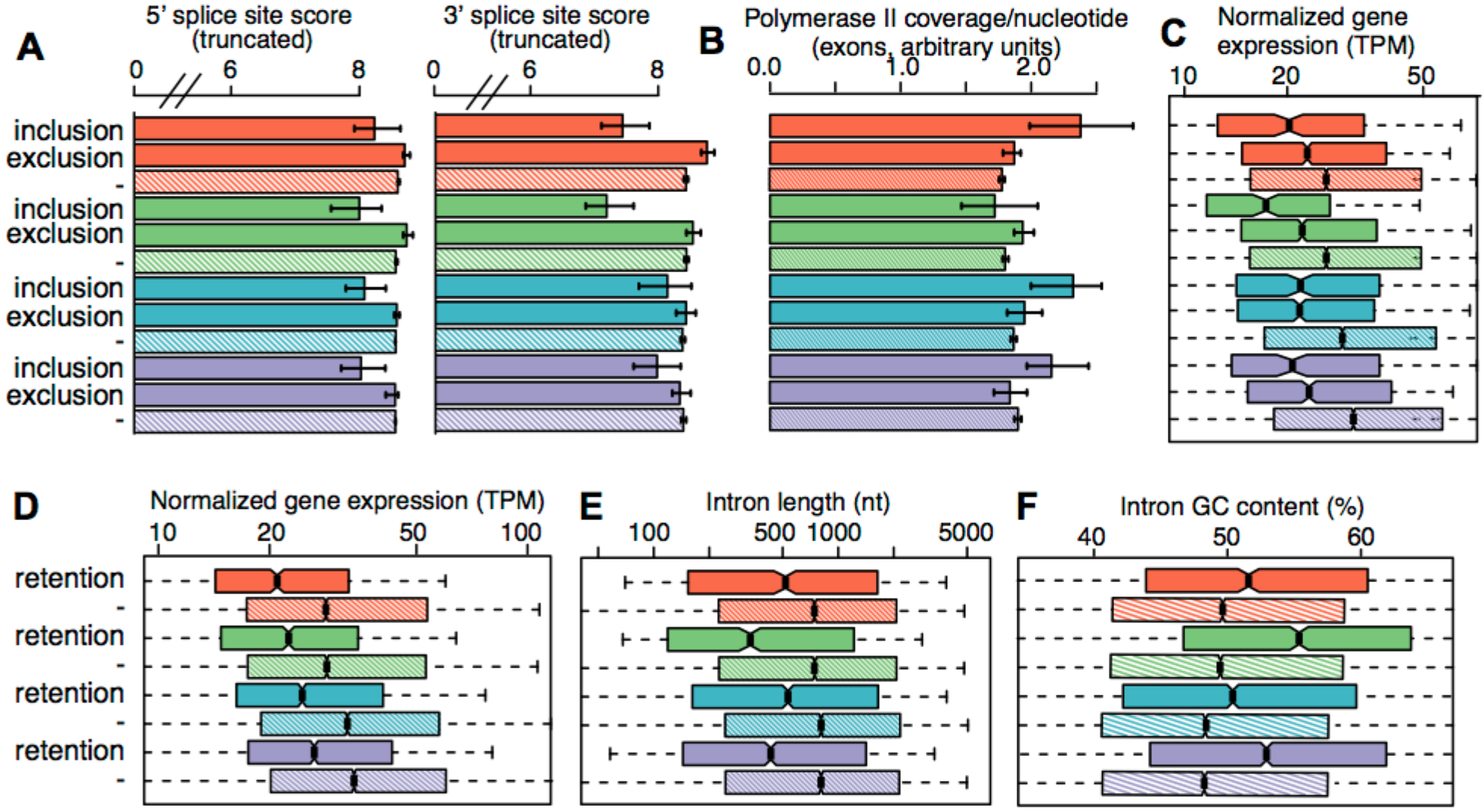
Susceptibility to snRNA depletion is associated with both *cis-* and *trans*-acting features. **(A)** Median scores of 5’ (left) and 3’ (right) splice sites, calculated with the MaxEnt method for the nucleotide sequences spanning the exon-intron boundaries (Yeo and Burge 2004), for exons with increased inclusion or exclusion following snRNA depletion.‘-’ indicates the genomic average (exons that are not differentially spliced, but which can be reliably detected within the illustrated sample). Error bars, 95% confidence intervals, estimated by bootstrapping. Colors as in Fig. 1A. **(B)** The average per-nucleotide RNA Pol II occupancy of alternatively spliced cassette exons. Error bars, 95% confidence intervals, estimated by bootstrapping. **(C)** Expression of genes containing differentially spliced exons and **(D)** introns. TPM, transcripts per million (TPM). Hinges represent the first and third quartile and notches illustrate the median +/−1.58 interquartile range scaled by the number of introns (IQR/√n), which approximately corresponds to the 95% confidence interval. Outliers not shown. **(E)** Distribution of intron lengths and **(F)** average GC content for introns that are retained following snRNA KD. indicates the genomic average (introns that are not differentially retained, but which can be reliably detected within the illustrated sample).

We next tested whether factors other than the splice site themselves might contribute to alternative exon usage following snRNA KD. As transcriptional rate has been previously demonstrated to influence cassette exon recognition (Fong et al. 2014; De La Mata et al. 2003), we wondered whether fast or slow transcription might similarly render specific exons sensitive or resistant to snRNA KD. We used RNA Polymerase II occupancy (Honkela et al. 2015) as an indicator of transcriptional rates: given constant levels of gene expression, reduced transcriptional speed indicates higher Pol II occupancy and vice versa. Cassette exons exhibiting increased inclusion following U1, U4, or U6 KD were characterized by increased Pol II occupancy relative to the genomic average as well as relative to cassette exons that were preferentially skipped following depletion of those snRNAs (**Fig. 4B**). While Pol II occupancy is an imperfect proxy for transcriptional speed, these results suggest that slower transcription of specific exons facilitates their recognition under conditions of lower snRNA abundance.

Increased Pol II occupancy of exons that were promoted by snRNA KD could, in principle, arise from increased gene expression (e.g., high density of fast-moving Pol II molecules) rather than slower transcriptional rates. We therefore tested for a relationship between gene expression and responsiveness to snRNA KD. Increased gene expression did not explain the increased Pol II occupancy of exons that were promoted by snRNA KD. Instead, these exons were preferentially located within genes exhibiting lower expression than genes containing exons whose inclusion was promoted by snRNA KD (**Fig. 4C**). Genes containing exons that responded to snRNA KD, whether with increased inclusion or exclusion, tended to be expressed at lower levels than the genomic average.

In contrast to cassette exons, introns whose recognition was affected by snRNA KD had neither weaker nor stronger 5’ and 3’ splice sites than the genomic average, suggesting that the initial recognition of exon/intron boundaries was not a determining factor for subsequent intron removal versus retention. Introns that were retained following snRNA KD were preferentially located within lowly-expressed genes (**Fig. 4D**). Differentially retained introns tended to be short and GC-rich. Introns that were retained following snRNA KD had a median length of 469 bp, versus 773 bp for introns that were not sensitive to snRNA KD (**Fig. 4E**). KD-responsive introns were approximately 1.9-5.9% more GC-rich than were unresponsive introns (**Fig. 4F**). Those trends are consistent with previous observations that intron length and GC content are strongly associated with an increased propensity towards intron retention (Dvinge and Bradley 2015). We next tested whether specific sequence motifs were enriched or depleted in KD-responsive introns by performing *ab initio* motif discovery. We recovered the expected signal for GC-rich sequence, but did not find other enriched or depleted sequence motifs that might be bound by known splicing enhancers or repressors.

### Breast cancer splicing profiles overlap with snRNA-modulated events

We next tested whether variable snRNA expression might contribute to the splicing dysregulation that characterizes most cancers. We observed a similar magnitude of variability in snRNA expression within our breast cancer cohort as we achieved via snRNA KD in our MCF7 model (**Fig. 1E**). We therefore hypothesized that the exons and introns that were differentially spliced following snRNA depletion *in vitro* (in MCF7 cells) would be preferentially mis-spliced *in vivo* (in breast cancer samples). To test this hypothesis, we performed deep RNA-seq for 136 of the invasive ductal carcinomas in our cohort and quantified global patterns of splicing. Anecdotal inspection of specific events, such as a retained intron within *LIME1*, revealed that biopsies with particularly low or high levels of a given snRNA frequently exhibited splicing patterns mimicking those observed following KD of that snRNA in MCF7 cells (**Fig. 5A**).

**Figure 5:**
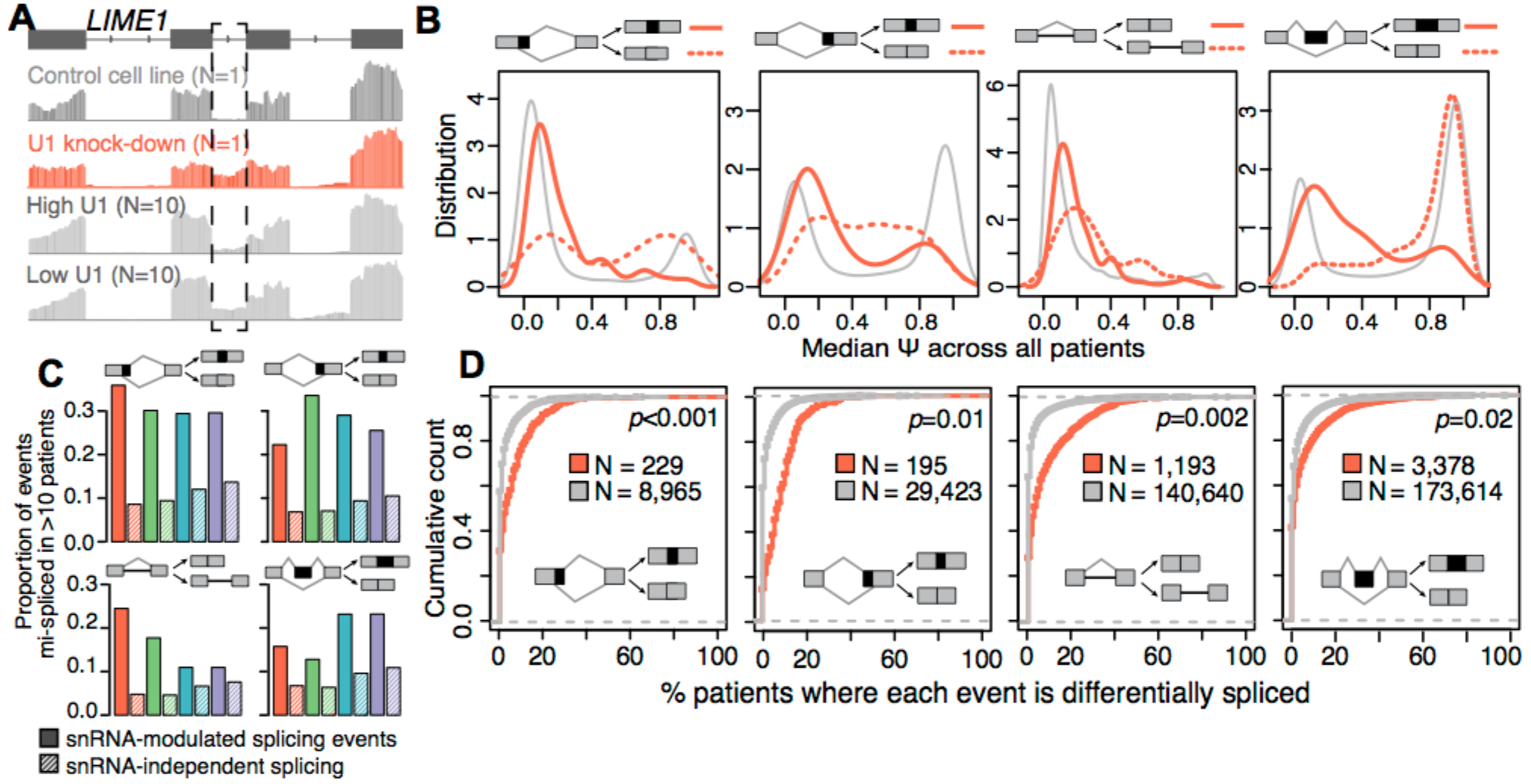
Dysregulated splicing in breast cancer is enriched for snRNA-responsive splice sites and exons. **(A)** snRNA-responsive intron excision within *LIME1.* Dashed box, differentially retained intron, for which U1 snRNA depletion promotes intron retention. Top rows, RNA-seq read coverage from control and U1-depleted MCF7 cells. Bottom rows, read coverage from the ten breast cancer biopsies with the highest and lowest U1 levels. **(B)** Distribution of Ψ values across breast cancer specimens (N = 136), stratified by U1 snRNA KD-sensitive (red) and -insensitive (grey) splicing in MCF7 cells. Dashed lines indicate splicing events for which the alternative spliced sequence is preferentially excluded from the mature transcript following U1 snRNA KD in MCF7 cells. Solid lines indicate events for which the alternatively spliced sequence is preferentially included following U1 snRNA KD. For retained introns, the dashed line indicates intron retention. Plots restricted to exons and introns that were annotated as alternatively spliced (see Methods). Note that events that are uniformly excluded or included from the mature mRNAs across the cohort have Ψ values close to 0 and 1, respectively, while intermediate Ψ values indicate heterogeneous splicing that varies within or across patients. **(C)** Proportions of snRNA-dependent and -independent events that are differentially spliced in at least 10 primary breast cancer biopsies relative to patient-matched peritumoral normal samples (N = 107). Samples are from the TCGA breast cancer cohort. Cumulative density functions comparing differential splicing of U1 snRNA KD-sensitive (red) and -insensitive (grey) splicing events across breast cancer patients, as in panel C. The plots illustrate the percentage of primary breast cancer biopsies exhibiting aberrant splicing relative to their patient-matched normal sample, across the TCGA cohort. Splicing events were stratified as U1 snRNA KD-sensitive (red) and -insensitive (grey) based upon the observed splicing in MCF7 cells. N, number of events that could be reliably quantified in the U1 snRNA KD sample. Statistical significance was calculated using the Kolmogorov-Smirnov test.

The biopsies that we characterized came from a genetically and clinically heterogeneous cohort, from which we intentionally selected samples representing multiple subtypes of cancer. Many factors in addition to snRNA levels influence the gene expression and splicing programs in each cancer; for example, different breast cancer subtypes exhibit markedly different levels of intron retention relative to peritumoral normal tissue (Dvinge and Bradley 2015). We therefore next tested whether snRNA-sensitive splicing events were frequently subject to consistent splicing dysregulation by measuring the uniformity of splicing for individual events across our entire cohort. For exons that were not affected by snRNA depletion, we observed a typical “on/off” splicing pattern, where the major isoform in almost all patients corresponded to near-complete inclusion or exclusion of the exon (**Fig. 5B**, left, exemplified by U1). In contrast, snRNA-responsive exons exhibited variable exon inclusion across the cohort, consistent with frequent alternative splicing/mis-splicing. We observed a similar pattern for retained introns as well as alternative 5’ and 3’ splice site events that responded to snRNA KD (**Fig. 5B**). We conclude that splice sites and exons that are particularly responsive to snRNA levels exhibit unusually variable recognition in different breast cancer patients, consistent with splicing dysregulation.

We next asked whether this unusually variable recognition of snRNA-responsive splice sites and exons corresponded to mis-splicing, in the sense of generating dysregulation relative to non-cancerous tissue. As matched normal control tissue was not available for our breast cancer cohort, we turned to data from The Cancer Genome Atlas, for which patient-matched breast cancer and peritumoral normal samples were available. We profiled genome-wide patterns of alternative splicing and identified the set of splicing events that were aberrantly spliced in ten or more patients relative to patient-matched normal tissue. These splicing events, which by definition were frequently dysregulated in cancer versus normal tissue, preferentially corresponded to events that responded strongly to snRNA KD *in vitro* (**Fig. 5C**). This enrichment was highly statistically significant; for example, competing 5’ and 3’ splice sites, retained introns, and cassette exons exhibited enrichments for U1 KD-responsive events with associated p-values of 0.001, 0.01, 0.002, and 0.02, respectively (**Fig. 5D**). Splicing events that were frequently mis-spliced in primary tumors relative to matched normal tissues exhibited highly similar and significant enrichments for U2, U4, and U6 KD-responsive events.

## DISCUSSION

Our data indicate that snRNAs play an unexpectedly complex role in establishing global splicing programs, in addition to their well-characterized roles in basal splicing catalysis. As perturbation of snRNA levels with physiological ranges induced widespread differences in alternative splicing, we suggest that snRNAs should be considered to act as regulatory in addition to basal factors.

In this manuscript, we focused on demonstrating that variable snRNA levels influence splice site and exon recognition, particularly in the context of breast cancer. However, our characterization of snRNA levels across multiple biological axes (**Fig. 1**) clearly demonstrates that snRNA levels are equally variable between tissues and developmental time points as they are between breast cancer patients. It is likely that snRNA levels play important and as-yet-unrecognized roles in establishing tissue-specific and developmental stage-specific splicing programs.

While we identified both *cis*- and *trans*-associated features that correlated with responsiveness to snRNA KD, the fundamental mechanisms underlying snRNA-mediated changes in splicing remain unknown. None of the features that we identified were strongly predictive of the observed splicing changes. Further work is required to determine why some splice sites and exons are so sensitive to altered snRNA abundance, while most splice sites and exons are resistant to perturbation of snRNA levels within physiological ranges.

Because we observed differential splicing following snRNA KD that was consistent with the known roles of each snRNA in basal splicing catalysis (**Fig. 2**), we speculate that each snRNA affects alternative splicing through distinct mechanisms. While KD of each snRNA resulted in a unique splicing profile, U1 and U2 KD preferentially impacted exon inclusion, whereas U4 and U6 KD preferentially impacted intron retention. These tendencies are consistent with the early requirement for U1 and U2 snRNPs’ interactions with the pre-mRNA. In contrast, as U4 and U6 snRNAs basepair with one another in the U4/U6.U5 tri-snRNP prior to spliceosome formation on the pre-mRNA, alterations in the relative abundance of those snRNAs might impede or promote the assembly of the pre-catalytic spliceosomal B complex and subsequent splicing catalysis.

As our data indicates that snRNA dysregulation shapes the global transcriptome of breast cancer, we speculate that snRNA dysregulation may contribute to tumorigenesis itself. Multiple prior studies have identified tantalizing connections between snRNAs and cancer. For example, overexpression of U1 snRNA may promote pro-cancer gene expression (Cheng et al. 2017), and U2 snRNA fragments are potential blood-based biomarkers for multiple cancers (Baraniskin et al. 2016; Kuhlmann et al. 2014; Köhler et al. 2016; Kuhlmann et al. 2015). Intriguingly, we observed a clear association between snRNA levels and the intrinsic breast cancer subtypes (Sorlie et al. 2001). The Her2 subtype displayed high concurrent levels of U1 and U5A, while the two clusters of triple-negative samples exhibited higher relative abundance of U6 or comparatively low levels of U2 and U5A. Further work is required to determine whether these differences in relative snRNA abundance are downstream of key regulators of each subtype, such as the estrogen or progesterone receptors, or instead contribute to the establishment of the gene expression programs that define each subtype. Regardless, it is clear that a full understanding of splicing dysregulation in cancer requires study of the RNA as well as protein components of the splicing machinery.

## METHODS

### Primary breast cancer sample collection and processing

Primary breast specimens were collected by the Fred Hutchinson Cancer Research Center/University of Washington Breast Specimen Repository with approval of the local IRB. Women were diagnosed with invasive breast carcinoma between 2002 and 2015 and had no prior cancer diagnosis and no neoadjuvant treatment (radiation, chemotherapy or hormone) prior to tissue collection. Breast tissue taken as core biopsies or surgical specimens were flash frozen and embedded in OCT. Each biopsy was macro-dissected into 10-20 10μm sections to enrich for tumor cells. Total RNA and DNA was isolated concurrently across sections from each biopsy, using the AllPrep DNA/RNA/miRNA Universal kit (Qiagen) according to manufacturer’s recommended protocol. IHC results were taken from medical records. Additional Ki67 testing was performed by immunohistochemistry. Tumor subtypes were determined using IHC results for estrogen receptor (ER), progesterone receptor (PR), HER2, and Ki67. The subtypes were defined as follows: Luminal A (ER and/or PR positive, HER2 negative, Ki67<14%); Luminal B (ER and/or PR positive, HER2 negative, Ki67>=14%); Her2 (Her2 positive, any ER/PR/Ki67); triple-negative (ER, PR, and HER2 all negative, any Ki67).

### Tissue specimens from healthy individuals

Total RNA from adult and fetal human tissues was obtained from Agilent Technologies. Adult tissue samples originate from individual donors, with the exception of breast, cerebellum, larynx, liver, lung, spleen, stomach, and trachea, which were from a pool of two to six donors each. Both genders were represented, with the age ranging from 56+/−16.5 years (median+/−standard deviation). All fetal samples were from donors aged 18 to 23 weeks. Brain, lung, and skeletal muscle were from individual donors, and the remaining fetal samples were pooled across two to 17 donors, aged 18 to 23 weeks.

### Cell culture and snRNA knock-down

MCF7 breast cancer cells (from American Type Culture Collection; HTB22) and HeLa cervical cancer cells (gift from J Cooper) were maintained in DMEM (4.5g/L glucose and glutamine, Life Technologies) with 10% fetal bovine serum, and 1% penicillin/streptomycin. MCF7 cell media was supplemented with 10μg/mL human recombinant insulin (Life Technologies). snRNA knockdown was carried out using chemically modified RNA-DNA hybrids (Integrated DNA Technologies), with a non-targeting scrambled sequence as control (Supplementary Table 2). Transfection was performed in six-well plates using Lipofectamine RNAiMAX (Invitrogen) following the manufacturer’s cell line-specific protocols (MCF7 reverse transfection; HeLa forward transfection), with 200pmole oligo and 10μL RNAiMAX reagent per well. Cells were harvested after 48 hours. Total RNA was extracted using NucleoSpin miRNA (Macherey-Nagel) to collect the small and large RNA fraction combined.

### snRNA quantification

snRNA levels in MCF7 and HeLa were quantified with real-time qPCR using the VeriQuest SYBR Green One-Step method (Thermo Fisher Scientific), with 100pg/μL totalRNA and 10μM primer per reaction (Supplementary Table 2). Samples were processed using the ΔΔCt method, using RN7SK as a reference gene, the knock-down with scrambled control as a calibrator sample, and primer-specific amplification efficiencies estimated from a four-sample dilution series. snRNA levels from human tissues and breast cancer specimens were quantified using the BioMark HD 48.48 Dynamic Array (Fluidigm Corporation). TotalRNA (5ng/μL) was mixed with 20X Sample Loading Buffer (Fluidigm Corporation), Fast EvaGreen qPCR Master Mix Lo-ROX (Biotium), Reverse Transcriptase and RiboSafe RNase Inhibitor (both SensiFAST One-Step, Bioline). Arrays were primed, loaded, and run according to the instrument specifications. All samples were run in triplicate, and were processed using the ΔCt method, using the median Ct value across RN7SK, RN7SL1 and 5S rRNA as a pseudo-reference gene, to correct for variations in amount of RNA input. ΔΔCt values were calculated relative to the median ΔCt values across the all tissues and the entire breast specimen cohort, respectively.

### RNA sequencing

RNA-seq libraries for cell lines samples were created with the KAPA Stranded mRNA-Seq Kit (Kapa Biosystems) with poly-A selection. Invasive breast carcinomas with a sufficient amount of high-quality totalRNA available were processed with the KAPA Stranded RNA-Seq Kits with RiboErase for ribosomal RNA depletion, due to potential mRNA fragmentation during storage of the specimens. Both types of libraries were prepared following the manufacturer’s instructions, with 500ng of total RNA as input, RNA fragmentation for 7min at 94C, and 10 PCR cycles of cDNA amplification. Library quality was assessed using the Agilent 2200 TapeStation. Barcoded RNA-seq libraries were sequenced in tri-plex on the Illumina HiSeq 2500 using single-ended 67bp reads.

### Genome annotations

Alternative splicing events were categorized as cassette exons, retained introns, and competing 5’ and 3’ splice sites, according to the MISO v2.0 annotations (Katz et al. 2010). Constitutively spliced junctions were defined as adjacent splice junctions where alternative splicing was not detected in any isoform of the UCSC knownGene track (Meyer et al. 2013). Separate annotation files were created for RNA transcripts and splice junctions for the read mapping. The RNA transcript annotation is a combination of isoforms present in MISO v2.0 (Katz et al. 2010) plus the UCSC knownGene (Meyer et al. 2013), and the Ensembl 71 gene annotation (Flicek et al. 2013). The annotation for RNA splice junctions contain an enumerating of all possible combinations of annotated splice sites as previously described(Hubert et al. 2013).

RNA-seq read mapping

The RNA-seq reads were mapped in five stages. 1) Bowtie (Langmead et al. 2009) and RSEM (Li and Dewey 2011) were used to map all reads to the UCSC hg19 (NCBI GRCh37) human genome assembly. Reads were mapped to the gene annotation file using RSEM with the arguments --bowtie-m 100 --bowtie-chunkmbs 500 --calc-ci --output-genome-bam after modifying RSEM v1.2.4 to call Bowtie v1.0.0 with the -v 2 mapping strategy. 2) The resulting BAM file was filtered to remove alignments that had a mapq score of 0 or where the splice junction overhang was 5bp or less. 3) Next, all the so far unaligned reads were extracted from the BAM file, and aligned to the RNA splice junction file, using TopHat v2.0.8b (Trapnell et al. 2009) with the arguments --bowtie1 --read-mismatches 3 --read-edit-dist 2 --no-mixed --no-discordant --min-anchor-length 6 --splice-mismatches 0 --min-intron-length 10 --max-intron-length 1000000 --min-isoform-fraction 0.0 --no-novel-juncs --no-novel-indels --raw-juncs. Other required parameters (--mate-inner-dist and --mate-std-dev) were determined for each sample by mapping reads to constitutive coding exons according to the exon_utils.py script in MISO. 4) The newly aligned reads were filtered again, using the same criteria as in stage 2. 5) Finally results from RSEM and TopHat were merged to create a combined BAM file containing all alignments. All transcriptome and splicing alignments were strand-specific.

### RNA splicing quantification

MISO (Katz et al. 2010) and v2.0 of its annotations were used to quantify splicing, using percent-spliced-in (PSI) values for all cassette exons, retained introns, and competing 5’ and 3’ splice sites. Reads directly spanning the splice junctions were used for detection and quantification of alternative splicing of constitutive junctions and retention of constitutive introns, as previously described(Hubert et al. 2013). All subsequent analyses were restricted to splicing events that were alternatively spliced in our data based on at least 20 relevant reads (*i.e.*, reads supporting either or both isoforms). For subsequently analysis of retained introns and alternative exon usage all statistically significant differentially spliced events were included, regardless of whether they were annotated as being alternative or constitutive.

### Sample clustering

Hierarchical clustering (heatmaps) of the relative snRNA levels within tissues and breast cancer patients was performed with the ‘ward.D2’ method, using data that had been standardized according to the ΔΔCt approach. MCF7 snRNA knock-down samples were clustered using multidimensional scaling (also known as principal coordinates analysis). The distances were calculated using the ‘canberra’ method, sum(|x_i_ − y_i_| / |x_i_ + y_i_|), using only events that were alternatively spliced in at least one sample, and had more than 20 reads supporting either of the spliced isoforms.

### Identification of differentially spliced isoforms

Events were defined as differentially spliced between a knock-down and the control if they satisfied the following criteria: 1) there were at least 20 relevant reads in both samples (reads supporting either or both isoforms), 2) there was a change in isoform ratio of at least 10%, and 3) there was a Bayes factor of statistical significance greater than or equal to 1. Wagenmakers’s framework (Wagenmakers et al. 2010) was used to compute Bayes factors for differences in splicing of individual events between sample pairs.

### Gene Ontology enrichment for differentially spliced events

Enrichment of Biological Process terms from the gene ontology was performed using the R package goseq (Young et al. 2010) using the “Wallenius” method. Splicing events were mapped back to genes, and compared to a background universe consisting of all spliced protein-coding genes with an expression level above 1 in at least two of the four knock-down samples, after normalizing the expression level within each sample using the trimmed mean of M values (TMM) method (Robinson and Oshlack 2010) with scaling factors calculated based on all protein-coding genes. The resulting false discovery rates were corrected using the Benjamini-Hochberg approach. Only terms with at least two ancestors were tested, to eliminate parent terms associated with generic biological processes.

### Transcription rates

RNA Polymerase II ChIP-seq and rRNA-depleted RNA sequencing data for MCF7 cells was obtained from GSE62789 (Honkela et al. 2015), using only the untreated samples. The RNA-seq data was processed as described above, with a maximum of 1 mismatch. The ChIP-seq reads were mapped to the human transcriptome using RSEM, following the same strategy. The gene expression values were normalized as above, and genes with expression <1 were filtered out, to reduce noise. For each individual exon the average per-nucleotide read coverage across the entire exon from start to end position was calculated.

### Analysis of breast cancer specimens from The Cancer Genome Atlas

TCGA RNA-seq data from breast cancer patient-matched tumors and samples from the adjacent normal tissue were obtained and processed as previously described (Dvinge and Bradley 2015) (N=107). To avoid bias due to events or genes predominantly expressed *in vitro* or *in vivo*, splicing events were filtered to include only events with a minimum of 20 event-specific reads, which could be detected in both MCF7 and at least 20 percent of patients. For the cumulative density function, the filtered events were stratified as snRNA KD-sensitive or -insensitive based upon the observed splicing in MCF7 cells. Statistical significance was determined using a one-sided Kolmogorov-Smirnov test, with the same number of steps as the cohort size (N=107).

### Data deposition

Sequencing data is deposited in the Gene Expression Omnibus (GEO) under accession number GSE107163.

## AUTHOR CONTRIBUTIONS

HD and RKB designed the experiments. JG and PLP identified and processed material from breast cancer biopsies. HD performed the experiments and created the figures. HD and RKB wrote the paper, with input from PLP and JG. All authors read and approved the final manuscript.

## ACKNOWLEDGEMENTS

This work was supported by the Seattle Tumor Translational Research program (HD) and Department of Defense Breast Cancer Research Program W81XWH-14-1-0044 (HD). RKB is a Scholar of The Leukemia & Lymphoma Society (1344-18). The results published here are in part based upon data generated by the TCGA Research Network: http://cancergenome.nih.gov/.

